# Statistical Learning of Distractor Suppression Down-regulates Pre-Stimulus Neural Excitability in Early Visual Cortex

**DOI:** 10.1101/2022.09.07.506943

**Authors:** Oscar Ferrante, Alexander Zhigalov, Clayton Hickey, Ole Jensen

## Abstract

Visual attention is highly influenced by past experiences. Recent behavioral research has shown that expectations about the spatial location of distractors within a search array are implicitly learned, with expected distractors becoming less interfering. Little is known about the neural mechanism supporting this form of statistical learning. Here we used magnetoencephalography (MEG) to measure human brain activity to test whether proactive mechanisms are involved in the statistical learning of distractor locations. Specifically, we used a new technique called rapid invisible frequency tagging (RIFT) to assess neural excitability in early visual cortex during statistical learning of distractor suppression, while concurrently investigating the modulation of posterior alpha-band activity (8-12 Hz). Male and female human participants performed a visual search task in which a target was occasionally presented alongside a color-singleton distractor. Unbeknown to the participants, the distracting stimuli were presented with different probabilities across the two hemifields. RIFT analysis showed that early visual cortex exhibited reduced neural excitability in the pre-stimulus interval at retinotopic locations associated with higher distractor probabilities. In contrast, we did not find any evidence of expectation-driven distractor suppression in alpha-band activity. These findings indicate that proactive mechanisms of attention are involved in predictive distractor suppression and that these mechanisms are associated with altered neural excitability in early visual cortex. Moreover, our findings indicate that RIFT and alpha-band activity might subtend different and possibly independent attentional mechanisms.

## Introduction

Adaptive human behavior relies on the ability to rapidly and efficiently select visual stimuli that provide useful, actionable information, and to ignore visual stimuli that are irrelevant. Statistical learning appears to play a prominent role in this kind of attentional control (Sherman, Graves, & Turk-Browne, 2020), and behavioral studies have shown that the spatial probability of distracting stimuli is implicitly learned and utilized to efficiently protect visual processing from interference (Ferrante et al., 2018; Sauter et al., 2018; Wang & Theeuwes, 2018; for a review, see Theeuwes et al., 2022). There are two possible mechanistic accounts for this effect of statistical learning on distractor processing. It may be that it is implemented proactively, with learned locations inhibited before stimuli appear. Alternatively, it may be that statistical learning of distractor suppression reflects the use of specific reactive inhibitory mechanisms that act after stimulus onset.

These alternatives are difficult to distinguish in overt behavior, and this has motivated the use of neurophysiological measures. For instance, behavioral indices of statistical learning of distractor suppression have correlates in the Pd event□related potential (ERP) component (Wang et al., 2019; van Moorselaar & Slagter, 2019), a post-stimulus evoked response closely linked to distractor inhibition (Hickey et al., 2009; Weaver et al., 2017). However, the results on Pd are inconsistent. In some cases, the Pd increases in response to distractors at high-probability locations (Wang et al., 2019), consistent with the idea that statistical learning supports stimulus-triggered distractor suppression. In other cases, the Pd decreases in response to distractors at high-probability locations (van Moorselaar & Slagter, 2019; van Moorselaar et al., 2020), and this has been interpreted as evidence that proactive suppression reduces the ability of distractors to draw selective resources, thus reducing the need for the stimulus-driven suppression reflected in the Pd.

If statistical learning of distractor suppression is implemented proactively, there are compelling reasons to expect this might be reflected in oscillatory neuronal activity in the alpha-band (8-12 Hz). Alpha power is strongly modulated by the allocation of spatial attention (Sauseng et al., 2005: Thut at al., 2006), is considered a neuronal marker of functional inhibition (Foxe & Snyder, 2011; Jensen & Mazaheri, 2010; Klimesch et al., 2007), and tracks strategic, cue-induced suppression of distractors (van Zoest et al., 2021). However, results investigating alpha’s role in statistical learning of distractor suppression are unclear, with some studies identifying increases in pre-stimulus alpha power over cortex contralateral to the most likely distractor location (Wang et al., 2019), while others finding no hemispheric lateralization (van Moorselaar & Slagter, 2019; van Moorselaar et al., 2020).

Pre-stimulus alpha may not be a consistent correlate of distractor suppression in all circumstances though (Foster & Awh, 2019, Noonan et al., 2016). Indeed, Gutteling et al. (2022) found that increased target-load in one hemifield was the main driver of the alpha power associated with a distractor in the opposite hemifield. Nevertheless, modulation of pre-stimulus alpha power by strategic suppression (for review, Chelazzi et al., 2019; Noonan et al., 2018) might differ from alpha modulation associated with statistical learning. Strategic suppression must be implemented quickly, whereas statistical learning necessarily evolves over time after many repeated experiences. Strategic distractor suppression is therefore likely to elicit prominent brain activity in the interval before distractors appear, reflecting rapid reconfiguration of neural systems, whereas statistical learning of distractor suppression could rely on latent neural representations that are not expressed in ongoing brain activity (Wolff et al., 2017; van Zoest et al., 2021). An alternative perspective is that it is not so much strategic versus implicit anticipation of distractors being the key modulator of the distractor related alpha. Recently, Gutteling et al. (2022) made the case the target load was the strongest modulator of the alpha associated with distractor suppression and they interpreted these findings by perceptual load theory. From this perspective, one would not predict a modulation of alpha power with statistical learning of distractors.

If pre-stimulus alpha is an ambiguous neural index of proactive distractor suppression in statistical learning, the pre-stimulus distractor suppression involved in this kind of visual learning might rely on a different neuronal mechanism. Here, we use magnetoencephalography (MEG) and a new technique called Rapid Invisible Frequency Tagging (RIFT; Hermann, 2001; Gulbinaite et al., 2010; Seijdel et al., 2022; Zhigalov et al., 2019; Zhigalov & Jensen, 2020) to assess neuronal excitability in visual cortex as statistical learning of distractor suppression emerges in behavior. If statistical learning of distractor suppression relies on proactive mechanisms, we expect this to emerge in the RIFT signal before the distractor appears.

## Materials and Methods

### Participants

Twenty healthy volunteers (13 females) took part in the study. One of the participants was excluded from the RIFT analysis due to a technical issue with recording equipment. All participants were included in the behavioral and time-frequency analysis.

The study was approved by the local ethics committee (University of Birmingham, UK) and written informed consent was acquired. Participants conformed to standard inclusion criteria for MEG studies, were right-handed, had normal or corrected-to-normal vision, and were financially compensated (£15 per hour).

### Experimental Design

We adopted a variant of the additional-singleton visual search task (Theeuwes, 1992; see Ferrante et al., 2018). Each trial began with a fixation point (500 ms), followed by presentation of four placeholder Gabor patches (6° x 6° visual angle, v.a.) presented at 4° v.a. from the fixation point for 1500 ms (Fig 1). At the end of this interval, a visual search array was presented for 300 ms. And subsequently replaced by a blank screen displayed until participant’s response. On distractor-absent trials (33% of the trials), the visual search array contained a target presented among three vertically-oriented non-targets of the same color. The stimuli were randomly either green or red. To ensure equivalent visual responses, the luminance of the two colors was matched using a flicker fusion procedure (Anstis & Cavanagh, 1983). On distractor-present trials (66% of the trials), the target was presented alongside a task-irrelevant distractor stimulus and two non-targets. This distractor consisted of a horizontally-oriented color-singleton patch, presented in the color that did not characterize the target and non-targets. Participants had to indicate the orientation of the oblique target patch (i.e., tilted either 15° to the left or to the right) within 2000 ms of the visual search array onset. They were asked to respond as fast and as accurately as possible. A new trial started after an inter-trial interval of 500 to 1500 ms (random selection from uniform distribution).

**Figure 1.**
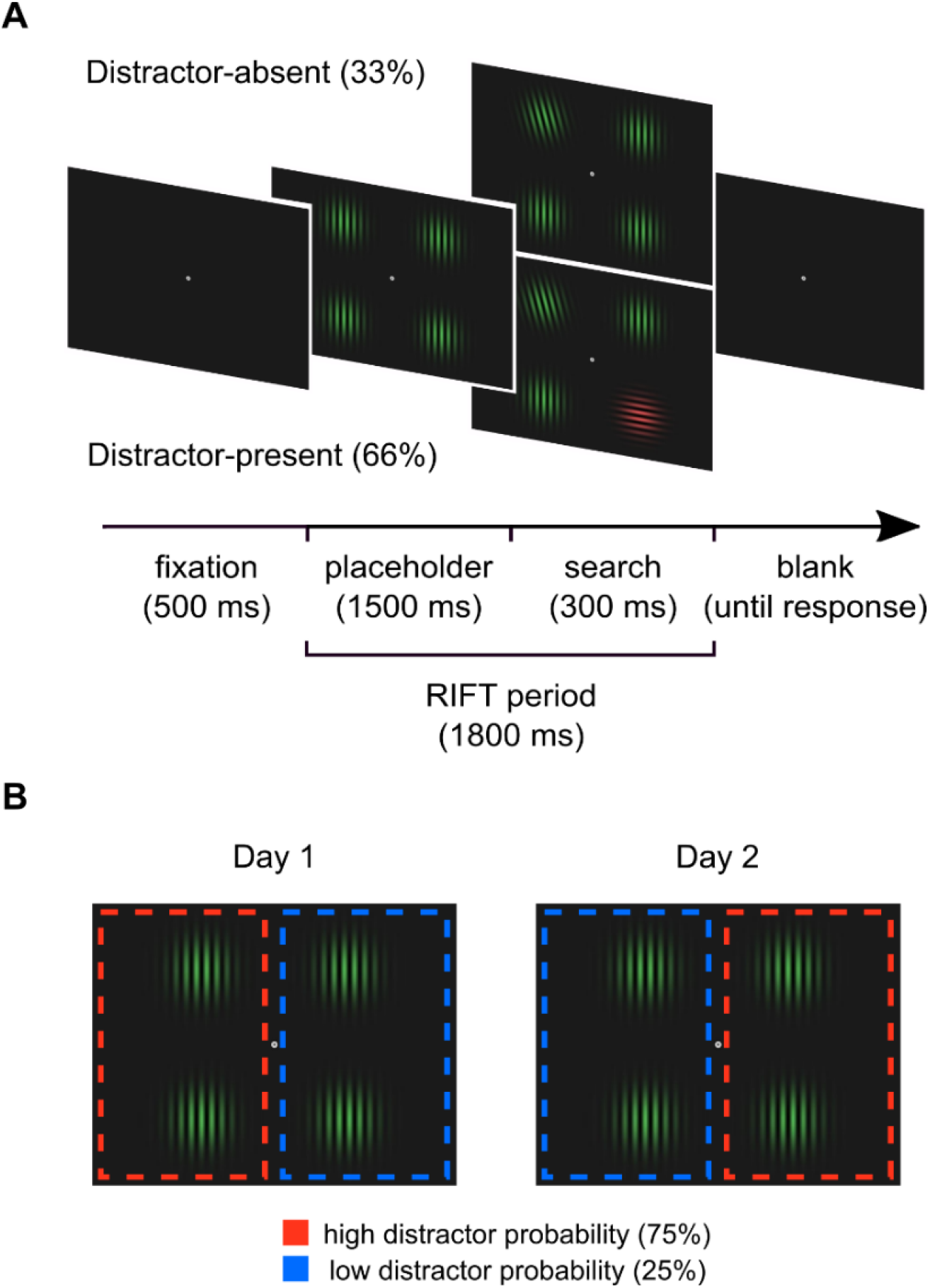
Experimental stimuli and procedure. A) The visual search task. Each trial started with a fixation dot. Afterwards, a placeholder screen with four Gabor patches was displayed. Then the search array was presented, and participants had to report whether the target was tilted to the left or right. In 66% of the trial, the target was presented together with a color-singleton distractor. During the whole RIFT periods, the two stimuli at the bottom were showed flickering at a broadband signal (RIFT) B) Statistical learning manipulation. The distractor was presented more often in one hemifield (75%) than the other (25%). To counterbalance the distractor probability conditions across the two hemifields, each participant took part in two experimental sessions (indicated as Day 1 and Day 2), with the statistical learning manipulation swapped after the first session.

To induce statistical learning of the distractor location, we biased the spatial probability of the distractors across the two hemifields (Fig. 1B). Specifically, the distractors were presented more frequently in one hemifield (75% of distractor-present trials; *high-probability hemifield*) than the other (25%; *low-probability hemifield*), while target probability remained equal across hemifields. Each participant completed two sessions of the experiment on two different days (indicated as Day 1 and Day 2 in Fig. 2B). Importantly, the statistical learning manipulation was swapped across hemifields after the first session to counterbalance the relationship between distractor probability and hemifield. The time gap between the two sessions was kept between 1 and 7 days and the initial assignment of the distractor probabilities to the two hemifields was counterbalanced across participants.

**Figure 2.**
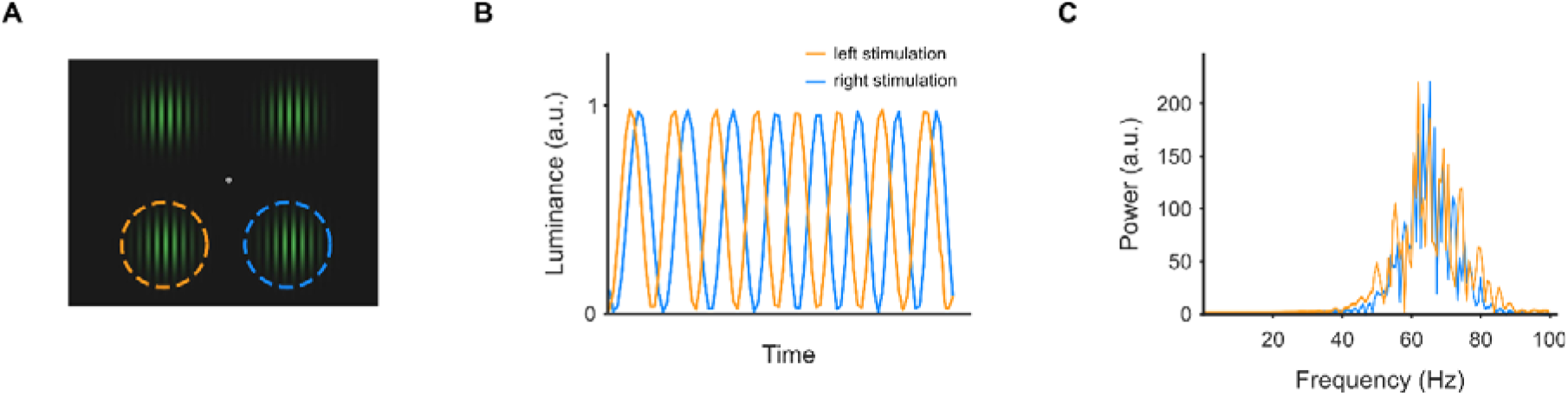
RIFT protocol. A) During the frequency tagging period, the two stimuli at the bottom of the screen (indicated by the dotted orange and blue circles) were flickered at very high frequencies (> 50 Hz) to generate a steady-state response in visual cortex. B) Examples of two uncorrelated (i.e., left and right stimulation signals) broadband tagging signals C) Power spectra of the broadband tagging signals used in this study.

Before starting the experiment, participants received verbal instructions and completed a practice block of 16 trials. The MEG sessions consisted of 6 blocks, each comprised of 10 mini-blocks. Each mini-block involved 12 trials in total: 4 distractor-absent and 8 distractor-present trials. In distractor-present trials, the distractor was presented 6 times in the high-probability hemifield (3 per location) and 2 times (25%) in the low-probability hemifield (1 per location). In total, each session consisted of 720 trials: 240 distractor-absent trials, 360 high distractor-present trials, and 120 low distractor-present trials. After the second session, participants filled out a debriefing questionnaire to evaluate whether they were aware of the statistical learning manipulation.

### Rapid invisible frequency tagging (RIFT)

RIFT involves the presentation of visual stimuli flickering at very high frequencies (> 50 Hz). Though this rate of flicker is not visible to participants, it generates a steady-state response in visual cortex that can be indexed in neurophysiological measures (Seijdel, Marshall & Drijvers, 2022; Zhigalov et al., 2019, Zhigalov & Jensen, 2020). Moreover, by employing independent flicker frequencies, RIFT can be used to measure neuronal excitability across different location representations in retinotopic visual cortex. This can be done without creating salient visual events at these locations and without interfering with endogenous oscillatory brain activity in the tagged frequency or in other frequency bands (Duecker et al., 2021). This allows us to concurrently a.) use RIFT to measure the impact of statistical learning in the pre-stimulus interval before stimuli appear, and b.) use time-frequency analysis to determine the role of alpha-band activity in modulating cortical excitability in this interval.

The interval of interest for RIFT analysis began with the onset of the placeholder screen and ended with the offset of the search array (see Fig. 1). In this time window, the luminance of the bottom-left and bottom-right stimuli (Fig. 2A) was modulated by two orthogonal broadband signals (Fig. 2B). To avoid discontinuity, the tagging signals were generated using frequency modulation as follows:

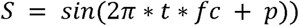

where *t* indicates the time variable (0 – 1.8 s), *f*_*c*_ is the frequency of the carrier signal (65 Hz), and *p* denotes the slow modulatory signal (i.e., noise with uniform distribution filtered within 2 – 6 Hz and scaled from -π to π). The spectral power of the tagging signals was limited to a frequency range of 55 to 75 Hz (Fig. 2C). To ascertain that the two tagging signals were independent of each other, each pair was generated by constraining the maximum correlation between the two tagging signals to 0.01.

### Projector

To achieve the rapid visual presentation required by RIFT, we employed a PROPixx DLP LED projector (VPixx Technologies Inc., Canada). This projector provides a refresh rate up to 480 Hz in color mode (and 1440 Hz in grayscale mode). This refresh rate is accomplished by dividing each frame received from the graphics card (at 120 Hz) into multiple frames. The projector divides each received frame (1920 × 1200 pixels) into four equally sized quadrants (960 × 600 pixels), allowing for a fourfold increase in refresh rate (480 Hz).

### MEG data acquisition and processing

MEG was acquired using a 306-sensor TRIUX MEGIN system at the Centre for Human Brain Health (CHBH). During acquisition, the MEG data were bandpass filtered between 0.1 and 300 Hz and sampled at 1000 Hz. The tagging signals were recorded using two custom-made photodiodes (Aalto NeuroImaging Centre, Finland) connected to the MEG system. Four head position indicator (HPI) coils were attached to the head of the participant to record the head position inside the helmet. A Polhemus Fastrack system (Polhemus Inc., USA) was used to save the location of HPI coils, as well as to digitize the fiducial points (nasion, left pre-auricular, right pre-auricular) and head shape. Electrocardiography (ECG) and electrooculography (vertical and horizontal EOG) were also recorded during the experimental session, and eye movements were recorded using an EyeLink 1000 infrared tracker (SR Research, Canada). Manual responses were collected using two five-button response boxes (NAtA Technologies, Canada).

MEG data were analyzed using MATLAB and the Fieldtrip toolbox (Oostenveld et al., 2011) following the standards defined in the FLUX Pipeline (Ferrante et al, 2022). All analyses reported here were conducted on gradiometer data. The data were detrended and segmented into 4 s epochs: -1.5 – 2.5 s relative to the onset of the tagging (placeholder screen). Epochs and sensors with artifacts were visually identified and excluded from further analysis. In order to focus our analysis on the time period in which statistical probability about distractor locations was established, we removed the initial 100 trials from the analyses. In addition, all error trials and trials in which participants responded within 200 ms from search array onset were excluded from analysis (mean 8.3% ± 4.1 SD), as were all trials where central fixation was not maintained (> 2.5° from central fixation point; mean 4.1% ± 5.9 SD). Independent Component Analysis (ICA) was used to identify blink- and cardiac-related components, which were removed from the data (mean 2.3 ± 0.8 SD).

### MRI data acquisition

High-resolution anatomical images were acquired for all participants with a 3-Tesla Siemens MAGNETOM Prisma scanner (T1-weighted MPRAGE; TR = 2000ms; TE = 2.01 ms; TI = 880 ms; flip angle = 8°; FOV = 256×256×208 mm; isotropic voxel = 1mm).

### Behavioral data analysis

Behavioral data analyses were performed using MATLAB and Jamovi (The Jamovi Project, 2020). Repeated-measures ANalysis Of VAriance (ANOVA) was performed on correct-trial reaction times (RTs) and accuracy, excluding trials in which the participants responded within 200 ms from search array onset (see below). We also removed the initial 100 trials from all analyses. When appropriate, p-values for statistical significance were adjusted for multiple comparisons using the Holm– Bonferroni method. Estimates of effect size are provided (η^2^_p_ or Cohen’s d) and non-significant results are accompanied by Bayes Factors (BF_10_ for t-tests and main effects in ANOVAs and BF_Inclusion_ for interactions in ANOVAs; Rouder et al., 2009).

### RIFT analysis

An analysis of the RIFT response based on the estimated power in the 55 – 75 Hz range would not be the optimal analysis as it would mix the broadband activity induced by the tagging stimulation with endogenous gamma activity. These two types of gamma-band activity have been shown to co-occur independently (Duecker et al., 2021). The cross-correlation analysis adopted here (described below), which exploit the time-dependent dynamic changes (phase and magnitude) in the tagging stimulation rather than the averaged power (magnitude only) in the frequency band, thus provide a better estimate of the neuronal excitability associated with visual input.

The RIFT response was estimated by computing the cross-correlation between the MEG signal and the tagging signal recorded through the photodiodes. Both MEG and photodiode recordings were first filtered between 55 and 75 Hz with a fourth-order Butterworth zero-phase filter. Pearson correlation was computed between each MEG sensor and photodiode, with these signals analyzed with a time lag between the signals starting from –200 ms offset and progressed in 1 ms steps to 200 ms offset. After applying the Hilbert transform, the cross-correlation indexes were normalized via min-max feature scaling and the hemifield-specific maximum correlation value estimated for each MEG channel (Zhigalov & Jensen, 2020).

To compare the tagging response to the high- and low-probability hemifields, we first computed the within-session difference between the left and right hemifields for each participant. Given the known focality of the RIFT effect (Zhigalov & Jensen, 2020), we identified the MEG sensor which showed the strongest correlation with the contralateral hemifield independently for each participant and session. Finally, we averaged the probability-specific tagging responses across sessions (i.e., hemispheres) independently for each participant.

### Time-frequency representations and alpha-band analysis

Time-frequency representations were calculated using a sliding time-window approach applied to the single trials. We used a 1000 ms time-window moved in steps of 50 ms. Each time frame was multiplied with a Hanning window prior to calculating the discrete Fourier transform (1 to 100 Hz in steps of 1 Hz). The power was then estimated as the modulus square of the Fourier transform. The power-estimates for each trial were averaged and a baseline correction was applied from -1 – - 0.5 s interval before the placeholder screen onset. To test for experiment-induced alpha activity lateralization, we computed a hemispheric lateralization index. First, the individual mean alpha power (8 – 12 Hz) in posterior sensors, separately for left and right hemifields, was averaged in the 0 – 1.5 s time window from placeholder screen onset. Then, the alpha lateralization index was obtained for each participant and each session by computing the difference between the mean posterior alpha power of the two hemifields.

### Source-localization analysis

To localize the source of both RIFT and alpha-activity, beamforming source-localization was performed using dynamical imaging of coherent sources (DICS; Gross et al., 2001) as implemented in Fieldtrip (Oostenveld et al., 2011). To build a forward model, we first manually aligned the MRI images to the digitized head shape. Then, the MRI images were segmented, and a single shell head model was prepared using surface spherical harmonics fitting the brain surface (Nolte, 2003). DICS spatial filters were computed for 10 mm grid where individual anatomy was warped into standard MNI template. Finally, the alpha lateralization index and the tagging response were computed for each grid point.

### Code accessibility

The experimental and analysis codes, as well as a copy of the debriefing questionnaire, are all available at https://github.com/oscfer88/dSL_RIFT.

## Results

### Statistical learning modulates distractor interference

We first analyzed raw RTs and accuracy data conducting a repeated-measures ANOVA with factors for Session (day 1 and day 2) and Distractor Location (distractor-absent, distractor in high- and distractor in low-probability hemifield). Participants responded faster during the second than the first session, F_(1,19)_ = 29.41, p < .001, η^2^_p_ = 0.61 (476 ± 11 ms vs. 515 ± 11 ms), while their performance did not differ in terms of accuracy, F_(1,19)_ = 0.53, p = .474, η ^2^ _p_= 0.03 (95 ± 0.9 % vs. 95 ± 0.9 %). Moreover, we found a significant main effect of Distractor Location on both RTs (F_(1,19)_ = 161.73, p < .001, η^2^_p_ = 0.90) and accuracy data (F_(1,19)_ = 4.60, p = .016, η^2^_p_ = 0.2).

To better understand the nature of the statistical learning effects, additional analyses were conducted on the behavioral cost produced by distractors presented at either high- or low-probability locations.

The cost of distractor presence on RTs was calculated for each participant as the difference between distractor-present and distractor-absent mean RTs. A repeated-measures ANOVA with factors for Session (day 1 and day 2) and Distractor Location (high- and low-probability hemifield) showed a significant main effect of Distractor Location, F_(1,19)_ = 21.76, p < .001, η ^2^ _p_= 0.53, confirming the effectivity of the statistical learning manipulation in modulating the interference produced by salient distracting stimuli. Indeed, distractor costs were larger when the distractor appeared in the low probability rather than high probability hemifield (mean ± SEM: 61 ± 4 ms vs. 48 ± 4 ms; Fig. 3A). The mean statistical learning effect was ∼13 ms (SEM: ± 3 ms; Fig. 3C). The main effect of Session was instead non-significant, F_(1,19)_ = 3.63, p = .072, η^2^_p_ = 0.16 Change in the statistical learning effect across sessions is captured in the interaction of Session and Distractor Location factors, and though this effect was not significant, F_(1,19)_ = 2.10, p = .163, η^2^_p_ = 0,1, BF_Inclusion_ = 1.17, the trend identified in this analysis warranted further consideration. To this end, we examined the statistical learning effect separately for each session. Distractor costs were significantly larger for low-probability than for high-probability conditions in session 1 and partially but not fully significantly in session 2, with the statistical learning effect emerging in both sessions with the expected directionality (Day 1: t_(19)_ = 4.02, p < .001, d = 0.9; low = 66 ± 4 ms, high = 48 ± 5 ms; Day 2: t_(19)_ = 1.78, p = .091, d = 0.4; low = 55 ± 5 ms, high = 48 ± 4 ms).

**Figure 3.**
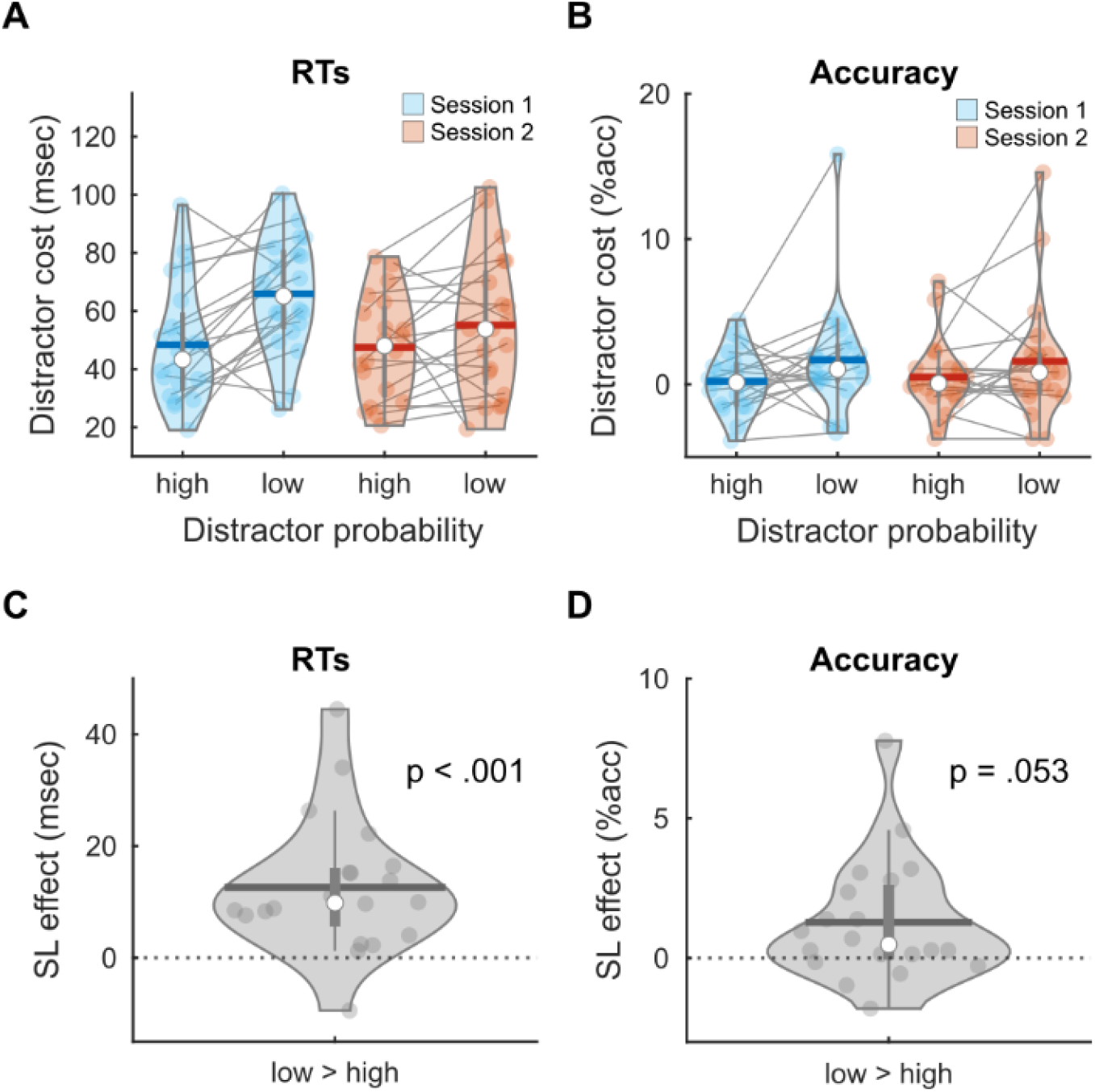
Behavioral results. Effect of distractor probability on distractor cost expressed in terms of A) reaction times (RTs) and B) accuracy divided by session. Overall, statistical learning (SL) effect on C) reaction times (RTs) and D) accuracy. The distractor cost was significantly smaller for high compared to low distractor probability in both RTs and accuracy. Blue dots representing individual median results, white dots representing averaged medians, blue horizontal lines representing averaged means, vertical thick line representing first (bottom edge) and third (top edge) quartiles.

Analysis of distractor costs on accuracy garnered similar results as the analysis of RTs. Namely, greater distractor interference emerged when the distractor appeared in a low-probability location (1.46 ± 0.72 % vs. 0.35 ± 0.42 %; Fig. 3B), which expressed as a marginally significant main effect of Distractor Probability, F_(1,19)_ = 4.25, p = .053, η^2^_p_ = 0.18. The average statistical learning effect was 1.11% (SEM: ± 0.54%; Fig. 3D). No other significant effects on accuracy were detected (Fs < 1).

To investigate how the learning progresses over time, we quantified the RTs and distractor costs blocked in 24 trials over time for high- and low-distractor conditions. From this analysis there was no clear evidence of a learning effect onset. This finding was consistent with extant literature suggesting that statistical learning is characterized by fast acquisition (see Ferrante et al., 2018; Jiang, et al., 2013).

Altogether, the behavioral results indicate that participants were able to learn the statistical probability of distractors and to respond faster and more accurately when distractors were presented in the more likely locations. Analysis of accuracy rules out the possibility that the RT effects were due to speed-accuracy tradeoff. Moreover, statistical learning emerged robustly in both RT and accuracy in each of the two experimental sessions. This allowed us to proceed with our plan to combine data from the two sessions for subsequent analysis.

### The RIFT signal reflects distractor probability

The analysis of the RIFT signal yielded a correlation peak when the cortical MEG signal lagged the visual signal by 60 ms, reflecting the time taken for the visual information to propagate from retinal ganglion cells through the lateral geniculate nucleus to primary visual cortex (Fig. 4A). Peak correlation emerged prominently in posterior sensors contralateral to the stimulation side (Fig. 4B). Source analysis localized the tagging response to early visual cortex (Fig. 4C).

**Figure 4.**
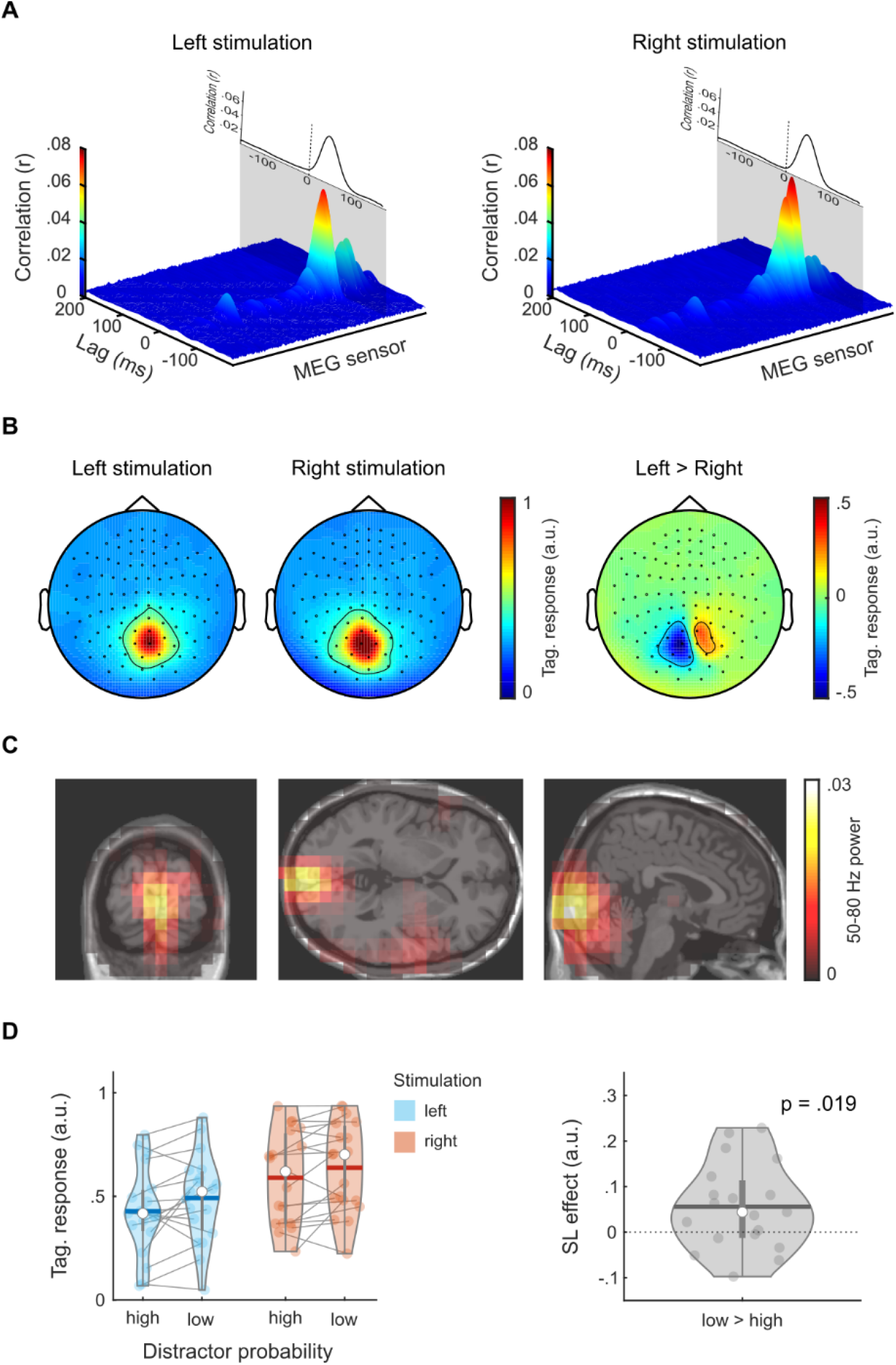
Rapid Invisible Frequency Tagging (RIFT) results. A) Results of the cross-correlation analysis for left and right stimulation across lags and MEG sensors (grand average). A correlation peak was detected at a time lag of ∼60 ms, reflecting the time taken for the visual information to reach the primary visual cortex. B) Sensor-level topological representation of normalized correlation value at peak lag for left and right stimulation and for their difference. The strongest correlation was observed over posterior-central sensors contralateral to the stimulated hemifield. C) Source-localization of the tagging response. The tagging response was localized in early visual cortex (V1/V2). D) Statistical learning effect on RIFT. The tagging response was significantly weaker for high compared to low distractor probability. Colored dots representing individual median results, white dots representing averaged medians, colored horizontal lines representing averaged means, vertical thick line representing first (bottom edge) and third (top edge) quartiles.

To investigate the effect of statistical learning on the RIFT signal, we first compared individual left and right tagging and identified the sensors that showed the strongest stimulation-specific response for each participant and session (Fig. 4B). Given that the distractor probabilities were swapped across hemifields between the two experimental sessions, we were able to evaluate the individual tagging response to the high- and low-probability conditions independent of distractor location (for location-specific results, see Fig. 4D, left plot). A repeated-measures ANOVA with factors for Stimulation Side (left and right) and Distractor Location (high- and low-probability hemifield) showed a significant main effect of Distractor Location, F(1,18) = 6.70, p = .019, η^2^_p_ = 0.27, with weaker correlation between the tagging signal and brain activity when the eliciting stimulus was in the location where the distractor was more likely to appear (0.51 ± 0.03 vs. 0.57 ± 0.03; Fig. 4D). The main effect of Stimulation Side did not reach full significance, F_(1,18)_ = 3.66, p = .072, η^2^_p_ = 0.17. The two-way interaction was also not significant, F_(1,18)_ = 0.41, p = .533, η^2^_p_ = 0.02. The results indicate that excitability was indeed reduced in early visual cortex located contralateral to the high probability distractor location.

To assess how the SL effect on RIFT evolved over time, we analyzed the SL effect on RIFT over sessions. A repeated-measures ANOVA with factors for Session (day 1 and day 2) and Distractor Location (high- and low-probability hemifield) show a significant main effect of Distractor Location, F_(1,18)_ = 4.79, p = .042, η^2^_p_ = 0.21, while the main effect of Session were not significant, F_(1,18)_ = 3.78, p = .068, η^2^_p_ = 0.17. The two-way interaction was also non-significant, F_p (1,18)_ = 0.93, p = .347, η^2p^ = 0.05.

An additional analysis was performed by dividing each session into three equal epochs. Given that the overall RIFT analysis was conducted on concatenated data, for this analysis the cross-correlation was recomputed on a single trial basis. A repeated-measures ANOVA including Distractor Probability, Stimulation Side, and Epoch (first, middle, and last third of the trials) as factors indicated a significant main effect of Distractor Probability, F(1,18) = 5.39, p = .032, η2p = 0.23, replicating our previous finding. The main effect of Epoch was also significant, F(2,36) = 5.99, p = .006, η2p = 0.25, with overall lower RIFT responses in the initial part of the experiment. However, this effect did not interact with Distractor Probability, F(2,36) = 0.60, p = .553, η2p = 0.03. These findings indicate that the statistical learning effect on RIFT does not vary systematically across sessions or different parts of the experiment, supporting the idea of fast adaptation to new statistical properties.

Furthermore, we assessed whether participants with stronger effect in terms of reduced distractor costs to learned distractors also showed a stronger RIFT modulation. We conducted a correlation analysis between the statistical learning effect expressed in terms of RTs and RIFT (i.e., low minus high distractor probability). However, no robust effects were found (r = .027, p = .913).

### No evidence of distractor probability modulations on alpha activity

To test whether the statistical learning manipulation had an impact on lateralized alpha-band activity (8 – 12 Hz), we first computed the power change relative to baseline for each participant and distractor probability configuration (i.e., high-probability on the left hemifield and low-probability on the right hemifield, and vice versa; Fig. 5A). We visually identified the sensors that showed the strongest overall alpha power modulation across participants during the placeholder display (i.e., from -1500 ms to the onset of the visual search array) compared to the baseline interval (−2500 to -2000 ms). These posterior sensors (identified in Fig 5) showed a decrease in occipital alpha activity in preparation for the visual search array. Source modeling showed that the alpha activity modulation was localized in occipital cortex (Fig. 5B). To test the impact of statistical learning on alpha band modulation, we compared the power in sensors located contralateral to the high- and low-probability hemifields (for location-specific results, see Fig. 5C, left plot). A repeated-measures ANOVA with factors for Hemisphere (left and right) and Distractor Location (high- and low-probability hemifield) did not identify a significant difference between high- and low-probability hemifields, F_(1,18)_ = 2.39, p = .138, η^2^_p_ = 0.11 (1.25 ± 0.07 vs. 1.26 ± 0.07; Fig. 5C). Bayesian analysis provided no evidence for or against the null hypothesis (BF_10_ = 0.65). All other effects were not significant (p-values > 0.17).

**Figure 5.**
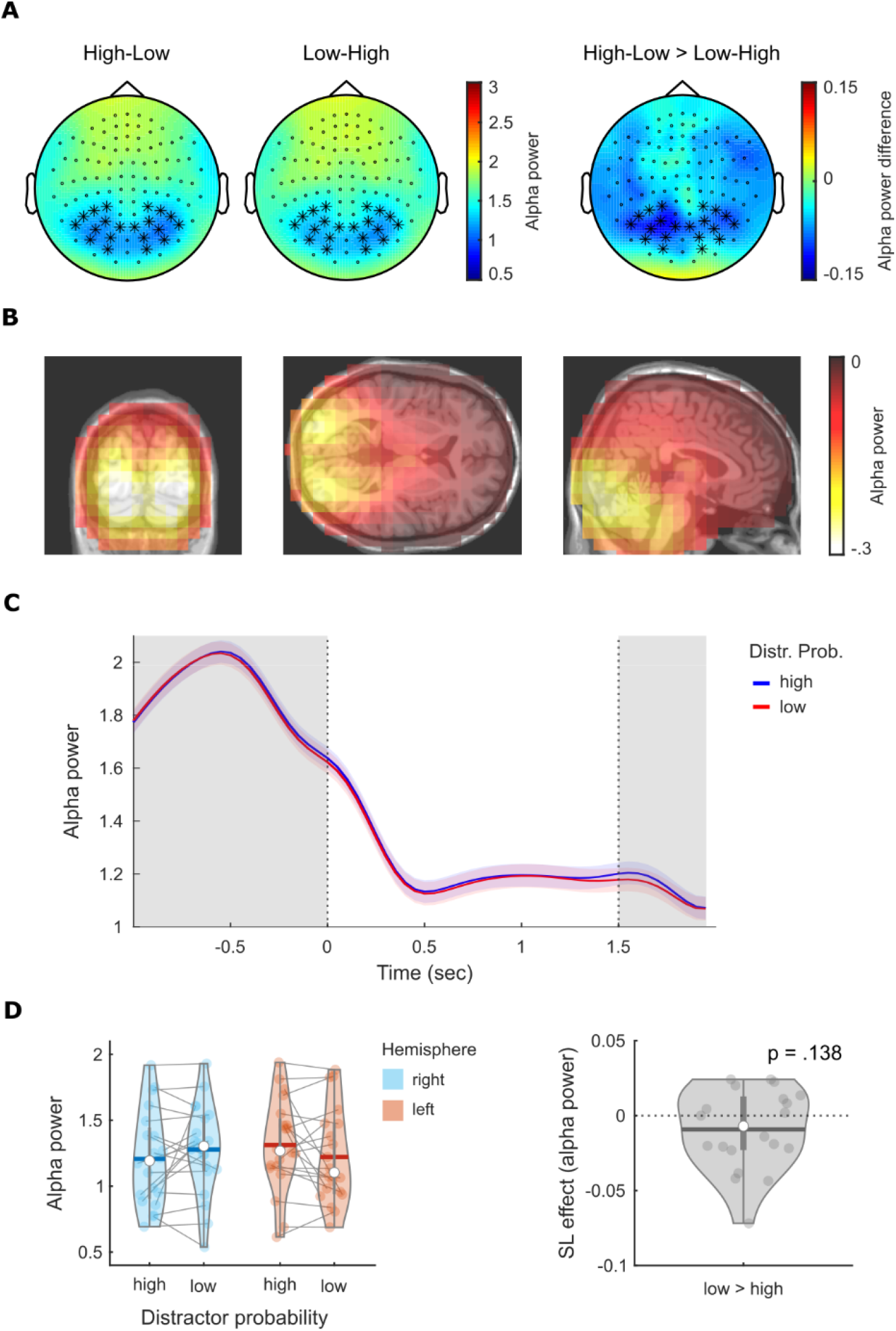
Alpha activity results. A) Sensor-level topological representation of baseline-corrected alpha power for the two different statistical leaning configurations and their difference. Asterixis indicates the sensors from where lateralized posterior alpha power was esteemed. B) Source-localization of alpha activity. C) Time course of alpha power by distractor probability. After the presentation of the placeholder display, contralateral alpha power decreased similarly for high- and low-probability condition. The left and right dotted lines represent the onset of the placeholder display and search array, respectively. D) Statistical learning effect on lateralized alpha power. There was no evidence for a modulation of distractor probability on alpha power.

Might the absence of a robust alpha lateralization be concealed by the baseline correction procedure we applied? To explore this possibility, we conducted a control analysis adopting the z-scoring method used in Wang et al. (2019) which allows to baseline-correct alpha power without subtracting pre-stimulus activity. To do so, we first computed the difference between posterior sensors located contralateral and ipsilateral to the high probability hemifield, separately for each trial. We then randomly swapped half of the trials and recomputed the difference (i.e., ipsilateral minus contralateral) 1000 times. This distribution was then used to transform the alpha lateralization index to z-scores for each participant. To account for possible *a priori* lateralization in alpha activity, this procedure was computed separately for each session. Sessions were then averaged within each participant before group analysis. The z-scored alpha lateralization index was averaged within the 0 - 1.5 s interval and tested against zero using the one-sample t-test. Results did not show any significant difference between (high-probability) contra- and ipsilateral alpha power (t(19) = 1.87, p = .767, d = .42), and Bayes analysis provided no appreciable evidence for or against the null hypothesis (BF10 = 1), thus replicating the previous null finding.

### Implicit nature of statistical learning

Fifteen of 20 participants completed the debriefing questionnaire. Three participants reported noticing that the spatial probability of the distractor was not equal across the four locations. One participant reported noticing that the spatial probability of the target was not equal across the four locations, though this was not the case. To further examine knowledge of the distractor probability manipulation, we asked participants to guess which side of the screen contained the distractor more often in the second session. Nine of the 15 correctly identified the higher probability location, χ^2^ (1, N = 15) = 0.6, p = .439. When they were asked to describe how confident they were in responding to this question, none of the 9 participants who correctly identified the high-probability location reported being “completely certain”, while 4 reported having “completely guessed”. These results suggest that participants were largely unaware of the spatial probability manipulation.

To assess whether participants who were relatively more aware of the statistical learning manipulation showed larger effects resulting in a more strategic form of suppression, we analyzed both RTs and accuracy data considering the reported awareness. The awareness level was based on the participant’s ability to correctly indicate the location of the high- and low-probability hemifields in the awareness questionnaire. Five participants were excluded from this analysis as they did not complete the questionnaire. Results did not yield any significant interaction between the distractor probability effect and the level of awareness (all p-values > .45). The same approach was used to the RIFT response and alpha power. Again, results did not show any significant interaction between the distractor probability effect and the level of awareness on either RIFT (all p-values > .80) and alpha power (all p-values > .48). Taken together, these results demonstrate that the level of awareness was unlikely to play a crucial role.

## Discussion

We tested the idea that statistical learning of distractor suppression reflects proactive suppression of neuronal activity associated with locations where distractors were likely to appear. We used a new approach called RIFT (Herrmann, 2001; Gulbinaite et al., 2019; Seijdel et al.,2022; Zhigalov et al., 2019; Zhigalov and Jensen, 2020) to assess neural excitability before the onset of target and distractor stimuli, showing that excitability in early visual cortex was significantly modulated by statistical learning of distractor suppression. In the interval before the onset of target and distractor stimuli, neural excitability was reduced in early visual cortex contralateral to locations where distractors were more likely to appear, consistent with the idea that statistical learning of distractor suppression emerges proactively, prior to stimulus onset. Notably, our results did not identify a relationship between this change in neural excitability and the emergence of posterior alpha-band oscillatory activity.

The RIFT approach relies on presentation of stimuli that vary in their contrast in a high-frequency band (>50 Hz) to produce steady-state visual evoked fields (SSVEF; Norcia et al., 2015; Vialatte et al., 2010). Importantly, participants are unable to detect the contrast changes employed in RIFT because they occur above the flicker fusion threshold, and the method accordingly provides the opportunity to measure neural excitability without the need for salient, visible events. Previous RIFT studies have revealed that the neural response in early visual cortex is enhanced contralateral to cued locations, reflecting the benefit of spatial attention on perceptual processing (Zhigalov et al., 2019). Here, we extend the technique by using RIFT to track visual prioritization when attention has not been explicitly cued.

Source modeling of the RIFT response observed in our study indicates that the neuronal alteration produced by statistical learning expresses in early visual cortex (V1/V2), in line with the effect of attentional cueing observed in earlier work (Zhigalov et al, 2019, Zhigalov & Jensen, 2020). This finding is also consistent with Zhang et al. (2021) who found regional signal suppression in early visual cortex using a similar statistical learning manipulation in fMRI. What drives this attentional modulation in early visual cortex? One possibility is that it reflects the action of fronto-parietal areas where visual priority of individual locations is topographically represented (Fecteau & Munoz, 2006; Ferrante et al., 2018; Zelinsky & Bisley, 2015). These “priority maps” are thought to contain a neural representation of the visual field that guides attention toward high priority locations, and this may involve suppression of information from low priority locations (Gaspelin & Luck, 2018).

An alternative is that the current results reflect predictive coding (Friston, 2005; Rao & Ballard, 1999). FMRI studies have demonstrated that expectations about the identity of a visual stimulus reduces pre-stimulus activity in primary visual cortex (Alink et al., 2010; Kok, Jehee & De Lange, 2012) and this has been interpreted as evidence that high-level neural populations convey predictions about upcoming events to lower-level areas, leading to a reduction of activity elicited by expected stimuli (Feldman & Friston, 2010). Foreknowledge of the distractor location in the current study may therefore have led to a reduction in neural excitability in cortex responsible for representation of distractors at this location.

Though the priority map and predictive coding accounts are impossible to distinguish in the current results, there is clear opportunity for further use of RIFT to differentiate between these hypotheses. For example, the idea that statistical learning is implemented in a fronto-parietal priority-map suggests that priority should be assigned to locations where the target occurs more often, leading to enhanced neural responses for stimuli at these locations. This contrasts with expectations from the predictive-coding framework, which suggests that the appearance of targets at predicted locations should generate a smaller neural response. The RIFT procedure has the potential to provide further perspective on the mechanisms underlying the change in cortical excitability identified in the current study.

The current results concretely demonstrate that statistical learning of distractor suppression is implemented proactively by modulating the neural excitability in early visual cortex. However, we were unable to unambiguously identify whether alpha activity plays a role in implementing this effect. This is in line with broader ambiguity on this issue in existing studies (van Moorselaar and Slagter, 2019; Wang et al. 2019). To date, one study has identified a reliable relationship between alpha and the statistical learning of distractor suppression, showing that lateral alpha increased in power contralateral to a high-probability distractor location, and this emerged only when analysis was limited to a low frequency band (7.5 - 10 Hz; Wang et al., 2019). Other attempts using powerful model-based approaches have garnered null results (van Moorselaar and Slagter, 2019; van Moorselaar et al., 2020). Here we found that alpha activity was broadly reduced in the pre-stimulus interval but did not find conclusive evidence either for or against the hypothesis that lateral alpha reliably tracked the manipulation of distractor probability.

The ambiguous alpha power results we see here, and that have emerged in earlier studies, may reflect a mixing of effects across participants or at different times within the experiment. The relationship between alpha oscillations and statistical learning of proactive distractor suppression is open to debate, but there is compelling evidence that the strategic implementation of proactive distractor suppression has a robust correlate in lateral alpha (van Zoest at al., 2021). If some participants were explicitly aware of the manipulation of distractor probability, or entertained this notion for periods in the experiment, our overall results might contain a lateral alpha power response that reflects strategic proactive distractor suppression over short period or for a minority of participants. While this kind of fleeting or inconsistent use of strategy in the sample would not be sufficient to generate the statistically robust RIFT and behavioral pattern we observe, it would make it difficult to statistically infer that contralateral pre-stimulus alpha was entirely unaffected by manipulation of distractor probability. This could underlie the inconclusive Bayesian statistics that we and others see in analysis of lateral alpha during statistical learning of distractor suppression. In addition, it might also be possible that the null finding on alpha is related to an issue on statistical power. However, even if this was the case, the RIFT results seem to indicate that the changes in excitability dominate the statistical learning of distractor suppression, with alpha power playing a secondary if any role.

If pre-stimulus lateral alpha activity proves to be a poor index of the statistical learning of distractor suppression, this could be the case for a few different reasons. As noted in the introduction, it may be that statistical learning of distractor suppression relies on ‘latent’ neural mechanisms instantiated in short-term synaptic plasticity or in tonic levels of neuromodulators. This kind of mechanism would not be directly evident in brain activity until ‘pinged’ by visual stimulation, like that employed in the RIFT technique, where the activity-silent modulation would express as a change in evoked neural activity. Activity-silent mechanisms of this kind have been identified in recent studies of working memory and are increasingly thought to play an important role in this context (Mongillo, Barak, & Tsodyks, 2008; Monohar et al., 2019). An alternative is that statistical learning impacts ongoing brain activity in the pre-stimulus interval, but that this precludes need for the kind of distractor suppression indexed in the alpha activity. Existing research has suggested that while the RIFT response indexes excitability in a circumscribed area in early visual cortex, alpha has a broader impact on higher-level representations in later visual areas (Gundlach et al., 2020; Gutteling et al., 2022; Peylo et al., 2021; Zhigalov & Jensen, 2020). If statistical learning expressed in a reduction of early cortical excitability, there may be no residual need for distractor suppression at the representational level impacted by alpha.

To conclude, the present study shows that information about the spatial probability of salient irrelevant stimuli is learned and used to better suppress distractors, with smaller behavioral interference found when distractors are presented at more likely locations. These attentional effects are reflected in a modulation of the RIFT response in early visual areas during the pre-stimulus period, indexing a relative reduction of neural excitability in the hemifield associated with higher distractor probability in advance of stimulus presentation. However, we did not find any evidence in favor of (not against) a possible involvement of posterior alpha activity. Statistical learning of distractor suppression thus proactively reduces the excitability and sensitivity of early visual cortex to visual stimulation.

